# Extracellular matrix protein anosmin-1 overexpression regulates dopaminergic phenotype in the CNS and the PNS with no pathogenic consequences in MPTP model of Parkinson’s disease

**DOI:** 10.1101/2022.11.25.517917

**Authors:** Javier Villadiego, Roberto García-Swinburn, Diego García-González, Rafael Lebrón-Galán, Verónica Murcia-Belmonte, Ernesto García-Roldán, Nela Suárez-Luna, Cristina Nombela, Miguel Marchena, Fernando de Castro, Juan José Toledo-Aral

## Abstract

The development and survival of dopaminergic neurons are influenced by the fibroblast growth factor (FGF) pathway. Anosmin-1 (A1) is an extracellular matrix protein that acts as a major regulator of this signaling pathway, controlling FGF diffusion, and receptor interaction and shuttling. Furthermore, overexpression of A1 *in vivo* gives rise to higher number of dopaminergic neurons in the olfactory bulb. Here, using A1 overexpressing mice (A1-mice), we studied the effects of A1 on different populations of catecholaminergic neurons in the central (CNS) and the peripheral nervous systems (PNS). A1 overexpression increases the number of dopaminergic SNpc neurons and alters the striosome/matrix organization of the striatum. Interestingly, these numerical and morphological changes in the nigrostriatal pathway of A1-mice do not confer an altered susceptibility to experimental MPTP-parkinsonism with respect to wild type controls. Moreover, the study of the effects of A1 overexpression was extended to different dopaminergic tissues associated with the PNS, detecting a significant reduction in the number of dopaminergic chemosensitive carotid body glomus cells in A1-mice. Overall, these analyses confirm A1 as a principal regulator of the FGF pathway in the development and survival of dopaminergic neurons in different nuclei of the mammalian nervous system.

## Introduction

Dopaminergic substantia nigra pars compacta (SNpc) neurons projecting to the striatum constitute the nigrostriatal pathway which, by the release of dopamine (DA) on their terminals, regulates the activity of striatal neurons participating mainly in the control of motor function (Albin et al. 1989; McGregor and Nelson 2019). The degeneration of this nigrostriatal pathway is the principal neuropathological feature of Parkinson’s disease (PD). In this neurodegenerative disease, the progressive neuronal death of SNpc neurons produces a significant DA deficit in the striatum that correlate with the characteristic motor symptoms (Braak et al. 2003; Obeso et al. 2017). In addition to the functional regulation of the striatal circuit by the release of DA, nigral neurons also participate in the embryonic formation of the striatal tissue since developing striosomes are established in the so-called “dopamine islands”. These “dopamine islands” are constituted by the incoming dopaminergic nigrostriatal fibers and are required for the proper striosome formation and differentiation (for review see: Crittenden and Graybiel 2011). Over the last years, significant advances have been developed in the knowledge of the molecular mechanisms by which SNpc neurons are specified, differentiated, and maintained (Arenas et al. 2015; La Manno et al. 2016). Among the different signaling pathways implicated in the development of SNpc neurons, fibroblast growth factor (FGF) pathway has shown to have a significant contribution for formation, differentiation, and survival of dopaminergic nigral neurons (Grothe et al. 2004; Timmer et al. 2007; Ratzka et al. 2012; Itoh and Ohta 2013). Despite the different morphogens implicated in the development of dopaminergic ventral midbrain neurons are relatively well identified, only few extracellular matrix components have been associated to this process (Gyllborg et al. 2018).

Anosmin-1 (A1) is an extracellular matrix glycoprotein that, among other effects, regulates FGF, Wnt, bone morphogenetic protein (BMP) and vascular endothelial growth factor (VEGF) signaling pathways during nervous system development (Endo et al. 2012; Korsensky and Ron 2016; Matsushima et al. 2020). A1 is encoded by the ANOS1 gene, former KAL1 (de Castro et al. 2017), whose mutation produces the Kallman syndrome (KS) that is characterized by hypogonadotropic hypogonadism and anosmia, presumably as a result of defects in neuronal migration and axonal targeting (Tsai and Gill 2006; de Castro et al. 2014). As an extracellular matrix protein, A1 interacts with different molecules of the extracellular space, including trophic factors and receptors, being the best studied up to date its interaction with the FGF receptor 1 (FGFR1) (Bülow and Hobert 2004; Murcia-Belmonte et al. 2010). Dual mechanisms have been described by which A1 regulates the FGFR1: *i)* in the named “productive interaction”, A1 binds with heparan-sulfate (HS) glycosaminoglycans and this complex acts as an amplifier of the signal produced by FGF2 to FGFR1; *ii)* in the so-called “abortive interaction”, if A1 concentrations exceed the amount of HS binding sites, free A1 can bind directly to FGFR1 inhibiting the receptor signaling (for review see: Korsensky and Ron 2016). In addition to these regulatory actions on FGFR1, A1 have also been identified as a key regulator for the FGFs diffusion (Wang et al. 2018). As a fundamental regulator of the FGF pathway, A1 has been shown to be essential for the formation of neural crest and sensory organs (Endo et al. 2012; Bae et al. 2018; Wang et al. 2018), to regulate the neurite branching (Soussi-Yanicostas et al. 2002; Gianola et al. 2009; Díaz-Balzac et al. 2015) and have been implicated in the migration and differentiation of neuronal and oligodendrocyte precursors (Bribián et al. 2006; García-González et al. 2010).

Despite the increased knowledge about the functional role of A1 in neural development obtained in the last decade by *in vitro* experiments or using different animal models (C. elegans, xenopus, zebrafish, and chick embryos), the lack of a mouse or rat orthologue for the ANOS1 gene has hampered an in-depth understanding of A1 functions in the mammalian nervous system. A transgenic mouse line that overexpresses the A1 human protein was generated some years ago in our laboratory, being the only mammalian model to study A1 functions *in vivo*. Using this A1 overexpressing mice (A1-mice) we demonstrated that A1 is involved in the subependymal zone (SEZ) neurogenesis and subsequent migration of neuroblasts towards the olfactory bulb (OB) during development, and the over-expression of A1 *in vivo* gives rise to significantly larger numbers of dopaminergic interneurons in the OB (García-González et al. 2016). In this work, we study the effect of A1 overexpression in the neuronal number and morphology of the principal dopaminergic nuclei of CNS. A1 overexpression produces a significant increase in the number of dopaminergic SNpc neurons, being similar as the raise previously described in the FGF2^-/-^ mice (Ratzka et al. 2012). Interestingly, A1-mice also show an altered organization of the striatal tissue, main target of projection of SNpc neurons. Moreover, we analyze the susceptibility of A1-mice to chronic 1-methyl-4-phenyl-1,2,3,6-tetrahydropyridine (MPTP) induced parkinsonism, showing a similar nigrostriatal degeneration than wild-type (wt) littermates. Finally, we study the effects of A1 overexpression on different dopaminergic tissues associated to the peripheral nervous system (PNS), finding that A1-mice show smaller carotid bodies with a reduced number of dopaminergic chemosensitive glomus cells. Our results indicate, by the first time, an important role of A1 in the genesis and maturation of SNpc dopaminergic neurons and in the determination of the striatal cytoarchitecture. In addition, the dissimilar effects of A1 overexpression found in the different dopaminergic nuclei analyzed suggest specifics niche-dependent actions of A1 in the development of dopaminergic neurons.

## MATERIALS AND METHODS

### Animals and chronic MPTP treatment

A1-mice and wt littermates were housed at regulated temperature (22±1ºC) on a 12 h light/dark cycle, with *ad libitum* access to food and water in the animal facilities of the Hospital Nacional de Parapléjicos (Toledo, Spain), the Instituto Cajal-CSIC (Madrid, Spain) and the CEA-Oscar Pintado, University of Seville (Seville, Spain). The generation of the transgenic mouse strain that over-expresses A1 was described previously (García-González et al. 2016; Murcia-Belmonte et al. 2016). In brief, regulatory elements of the human β-actin promoter carried by pBAP vector were used to direct the expression of the human ANOS1 cDNA, and a downstream IRES-EGFP cassette was used as a reporter. All the experiments were performed using 2-6 months old male mice in accordance with the European Communities Council Directive (2010/63/UE) and the Spanish (RD53/2013) regulations. Chronic MPTP treatment was carried out, as previously reported (Muñoz-Manchado et al. 2016; Villadiego et al. 2018a), by the subcutaneous (s.c.) administration of MPTP (20 mg/kg; Sigma) 3 times per week for 3 months. A control group received vehicle solution (0.9% NaCl; Sigma). Mice treated with MPTP, or vehicle solution, were killed 8 days after the end of the MPTP treatment, the brains were immediately removed and fixed as described below. Animals were sacrificed under deep anesthesia induced by a mix of 100 mg/ kg ketamine (Pfizer) and 10 mg/kg xylazine (Bayer). The animal procedures were approved by the Animal Review Board at the Hospital Nacional de Parapléjicos (registered agreement number SAPA001), Instituto Cajal (CEEA-IC), and the Animal Research Committee of the University Hospital Virgen del Rocío (University of Seville; registered agreement number 11-07-14-112).

### Histological analyses

Mice were transcardially perfused with 50 ml of PBS (Sigma) and 50 ml of 4% paraformaldehyde (Sigma) in PBS. Brains, carotid bifurcations, and suprarenal glands were removed and fixed overnight at 4ºC with 4% paraformaldehyde in PBS. After fixation, tissues were cryoprotected in 30% sucrose (Sigma) in PBS and included in Optimum Cutting Temperature compound (O.C.T. compound, Tissue-Tek). Tissue sections were sliced using a cryostat (CM3050-S; Leica). Brain coronal sections were cut at 30 μm of thickness, and 10 μm sections for the superior cervical ganglia (SCG), carotid body (CB) and suprarenal gland tissues. Immunohistological detection of tyrosine hydroxylase (TH; NB300-109, Novus Biologicals, RRID:AB_10077691, 1:1000), anti□adenomatous polyposis coli, clone CC1, (CC1; OP-80, Calbiochem, RRID:AB_2057371, 1:200) and glial fibrillary acid protein (GFAP; Z0334, Dako, RRID:AB_10013382, 1:500) were performed as previously described (Muñoz-Manchado et al. 2013; Ortega-Sáenz et al. 2013; Murcia-Belmonte et al. 2016), using a secondary peroxidase-conjugated antibody kit (K4011, Dako) or, for immunofluorescence, anti-rabbit IgG-Alexa 568, anti-mouse IgG-Alexa 488 secondary antibodies (respectively A11036, RRID:AB_10563566 and A21202, RRID:AB_141607, 1:400; Invitrogen) and 4’,6-diamidino-2-phenylindole (DAPI; 1:1000; Sigma) for nuclei counterstaining. DAB-stained slides were mounted on Glycerol: PBS-Azide 0.02% (1:1), and fluorescent slides were mounted on DAKO Fluorescence Medium (DAKO, S3023).

The eriochrome cyanine histological staining of myelin was performed following a previously described protocol (Kiernan 2007). Briefly, 30 μm brain coronal sections were mounted on Superfrost Plus™ adhesion slides (J1800AMNZ, Fisher Scientific) and dried at 37ºC for 30 minutes. After that, slides were placed in eriochrome cyanine solution (0,1% eriochrome cyanine R, 5% ferric ammonium and 0.25% sulfuric acid; Sigma) 30 minutes at r.t and washed with tap water. Then, the staining was differentiated with 10 minutes of incubation in 5% ferric ammonium sulphate solution (Sigma) and 10 minutes in borax-ferricyanide differentiator (1% sodium tetraborate and 1.25% potassium hexacyanoferrate; Sigma) at r.t. After 3 washes with tap water, the tissue was dehydrated with 70%, 95% and 100% EtOH, and twice in xylol (5 minutes at r.t for each step; Sigma). Finally, eriochrome-stained slides were mounted on LEICA CV mounting medium (#14046430011, Leica).

Images were acquired by a light-transmitted microscope (BX61, Olympus) equipped with a refrigerated digital camera (DP70, Olympus) or by a confocal microscope (DM IRE2, Leica).

### Stereology and densitometry

Unbiased stereological analysis was performed by systematic random sampling using the optical dissector method (West 1993) for the estimation of the number of TH^+^ neurons. Stereological estimations were carried out in the region spanning: (i) from - 2.80 mm to -3.28 mm for the substantia nigra pars compacta (SNpc) and the ventral tegmental area (VTA); (ii) from -5.34 mm to -5.80 mm for the locus coeruleus (LC); from 1.54 mm to -0.70 mm for the striatum (Str); (iii) from -1.06 mm to -2.80 mm for the zona incerta (ZI); and (iv) from -1.22 mm to -2.77 mm for the arcuate nucleus (Arc) relative to Bregma according to the Franklin and Paxinos mouse brain stereotaxic atlas (Paxinos and Franklin 1997). Reference volumes for each section were outlined under low magnification (4x), and TH^+^ neurons were counted at high magnification (40x) using the following optical dissectors: 7225 μm^2^ x 20 μm for SNpc and VTA TH^+^ neurons, and 6400 μm^2^ x 20 for LC, ZI and Arc; with a guard volume of 5 μm to avoid artefacts on the cut surface of the sections. Striosomes and striatal CC1^+^ cells density was analyzed using, respectively, 40000 μm^2^ and 44066 μm^2^ counting frames. Volume estimation was carried out using the Cavalieri principle (Gundersen and Jensen 1987; Howard and Reed 2010). TH^+^ neuronal soma volume and striosome area was determined using the nucleator (Gundersen 1988). TH^+^ and GFAP^+^ CB cells were determined as previously described (Ortega-Sáenz et al. 2013; Macías et al. 2014). All stereological procedures were performed using the appropriate tools of the New CAST™ system (Visiopharm) or NIH Image software (FIJI-ImageJ).

The optical density (O.D.) of striatal TH^+^ and CB GFAP^+^ staining was measured from digitized pictures using the NIH Image software (FIJI-ImageJ) as previously described (Muñoz-Manchado et al. 2016; Trillo-Contreras et al. 2018). Striatal TH ^+^ or CB GFAP^+^ O.D. values of each animal were obtained from a total of 6 slices covering the entire rostro-caudal extent of the same striatal region analyzed by unbiased stereology (see above) or 4-6 slices covering the whole CB tissue.

### HPLC

Striatal catecholamine content was measured by HPLC as previously reported (Villadiego et al. 2018b). Briefly, striatal tissue was obtained fresh by dissection in ice-cold PBS under a stereoscopic binocular microscope (Olympus SZX16). Samples were frozen immediately in liquid N_2_ and kept at -80ºC until its use. Striata were sonicated in 200 μl of chilled solution containing 0.1 M HClO_4_, 0.02% EDTA and 1% ethanol (Sigma) and centrifuged at 16000 g for 10 min at 4ºC. Supernatants were filtered with a 30000 Da molecular mass exclusion membrane (Millipore) by centrifugation at 16000 g for 30 min at 4ºC and injected into an HPLC system (ALEXYS 100; Antec Leyden). Dopamine (DA), 3,4-dihydroxyphenylacetic acid (DOPAC) and homovanillic acid (HVA) levels were determined using a 3 μm C-18 column (ALB-215; Antec Leyden) by electrochemical detection with a glassy carbon electrode and in situ ISAAC reference electrode (Antec Leyden). Concentrations of compounds were expressed as ng/mg of total protein. Pelleted proteins were resuspended in 0.1 M NaOH (Sigma) and assayed for protein quantification using the Bradford assay (Biorad).

### GDNF ELISA

GDNF protein content was measured from striatal tissue using the methodology described before by our group (Villadiego 2005; Rodriguez-Pallares et al. 2012). Briefly, striatal tissue was dissected as indicated before, frozen in liquid N_2_ and homogenized in lysis buffer (137mM NaCl; 20mM Tris, pH 8.0; 1% IGEPAL CA-630, 10% glycerol, 1/1000 protease inhibitor cocktail; Sigma) using a Polytron (OMNI). Protein extraction and the ELISA assay were performed following the manufacturer instructions, with the exception of the anti-GDNF monoclonal and anti-hGDNF polyclonal antibodies that were used at 1:500 and 1:250 respectively. Absorbance at 450 nm was measured in a plate reader (Thermo) and the total protein content of the samples was obtained by the Bradford assay (Biorad). As GDNF is not expressed in the mouse cortex, cortical protein lysates were used to determine the ELISA background signal (Enterría[Morales et al. 2020).

### Statistical analysis

The specific number of mice or cells analyzed (n) on each experimental group is indicated in each figure or figure legend. Data are presented as mean ± SEM. In all cases, the normality test (Kolmogorov-Smirnov) and the equal variance test were carried out, and, when passed, the Student’s t-test. In the cases that the normality test failed, the non-parametric Mann–Whitney U test was performed. All statistical analyses were conducted using Sigmastat 2.0 (RRID:SCR_010285) or Graphpad Prism7 (RRID:SCR_002798) software.

## RESULTS

### Anosmin-1 overexpression produces higher number of SNpc dopaminergic neurons

In a previous study it was reported that A1 overexpression induces an increase in the number of TH^+^ interneurons in the olfactory bulb (García-González et al. 2016). Here, using the same transgenic mouse line, we investigated how the increased expression of A1 could alter the number and the morphology of neurons from a principal CNS dopaminergic pathway, the nigrostriatal pathway. The general aspect of the substantia nigra pars compacta (SNpc) was not altered in the A1 overexpressing mice respect to their wt littermates (Fig. 1A). However, the stereological quantification of the SNpc TH^+^ neuronal number revealed a significantly augmented number of dopaminergic neurons in A1-mice respect to wt controls (Fig. 1B). In order to study if this increased number of dopaminergic SNpc neurons found in A1-mice is due to a general effect of A1 overexpression on CNS catecholaminergic neurons or, on the contrary, is related with specific actions of A1 on certain TH^+^ nuclei, we examined the TH^+^ neuronal number of other CNS catecholaminergic nuclei as the locus coeruleus (LC), ventral tegmental area (VTA), zona incerta (ZI) and arcuate nucleus (Arc). Almost 50% more TH^+^ neurons were also found in the LC of A1-mice in comparison to wt-mice (Fig. 1C-D), while the number of dopaminergic neurons in other TH^+^ nuclei (VTA, ZI and Arc) did not change with the A1 overexpression (Fig. 1E). In addition to the TH^+^ neuronal number quantification, we performed a stereological analysis of the soma volume of TH^+^ neurons in all these structures. Interestingly, we observed a generalized soma volume decrease of these TH^+^ populations in A1-mice, varying from ∼40% of reduction for LC neurons to just a trend of ∼10% of decrease in the case of ZI and Arc neurons (Fig. 2). Altogether, our morphological analyses showed that the *in vivo* overexpression of A1 produces a ubiquitous shrinking of the neuronal soma volume of different CNS catecholaminergic neurons and an increased number of TH^+^ neurons that is restricted to specific regions such as SNpc and the previously described olfactory bulb (García-González et al. 2016), and the LC with a clear trend.

**Figure 1.**
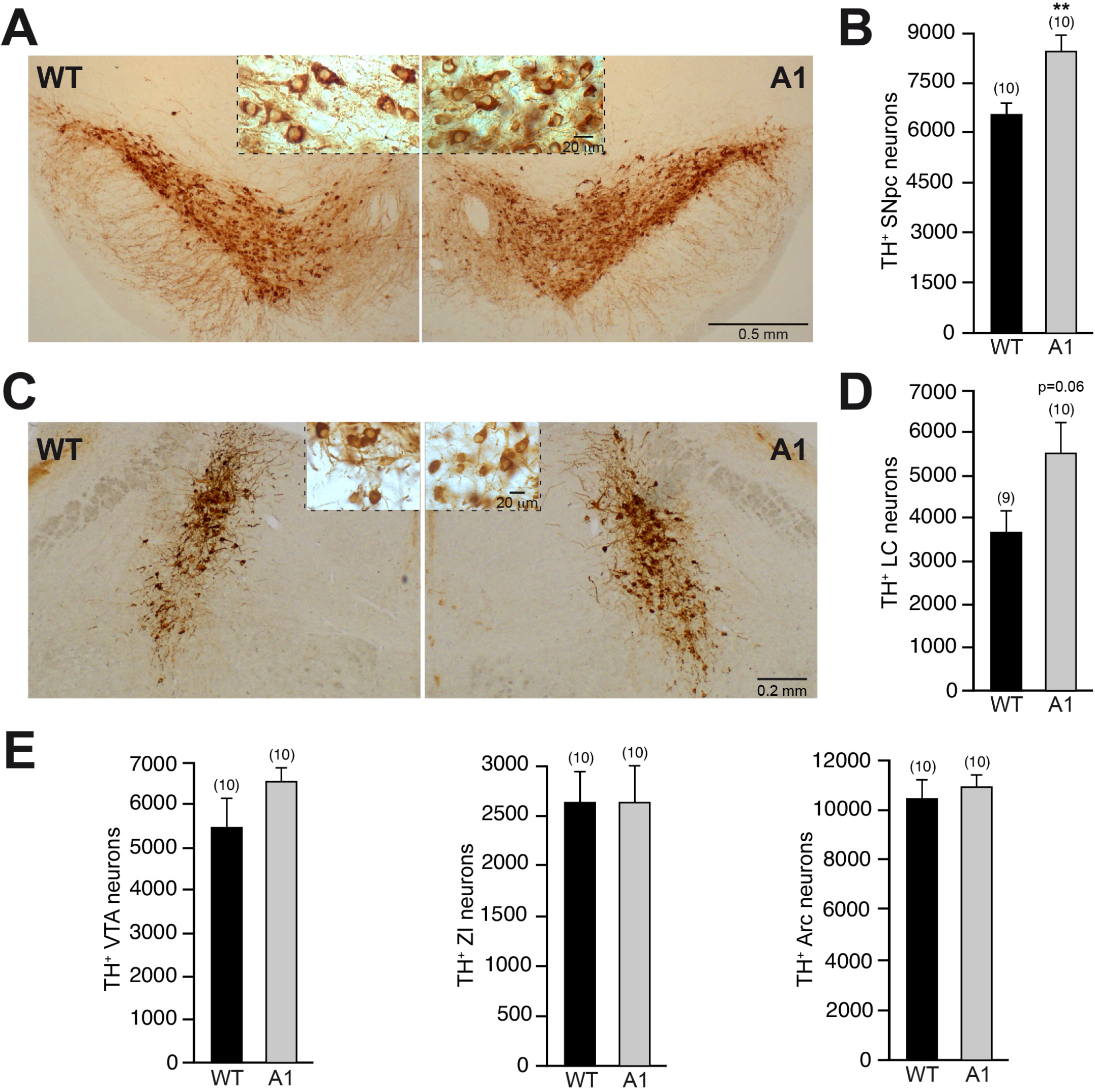
Analysis of TH^+^ neurons in different CNS catecholaminergic nuclei of A1-mice. A-B: Representative images of mesencephalic coronal sections, after TH immunohistochemistry, from wt and A1-mice. B: Stereological quantification of SNpc TH^+^ neurons. C: Representative images showing catecholaminergic neurons of LC from wt and A1-mice. D: Stereological quantification. E: Quantification of the number of TH^+^ neurons in the VTA, ZI and Arc from the previously described experimental groups. Data are presented as mean ± standard error of the mean (S.E.M.). The number of mice analyzed are shown between brackets. Unpaired t-test. **p<0.01.

**Figure 2.**
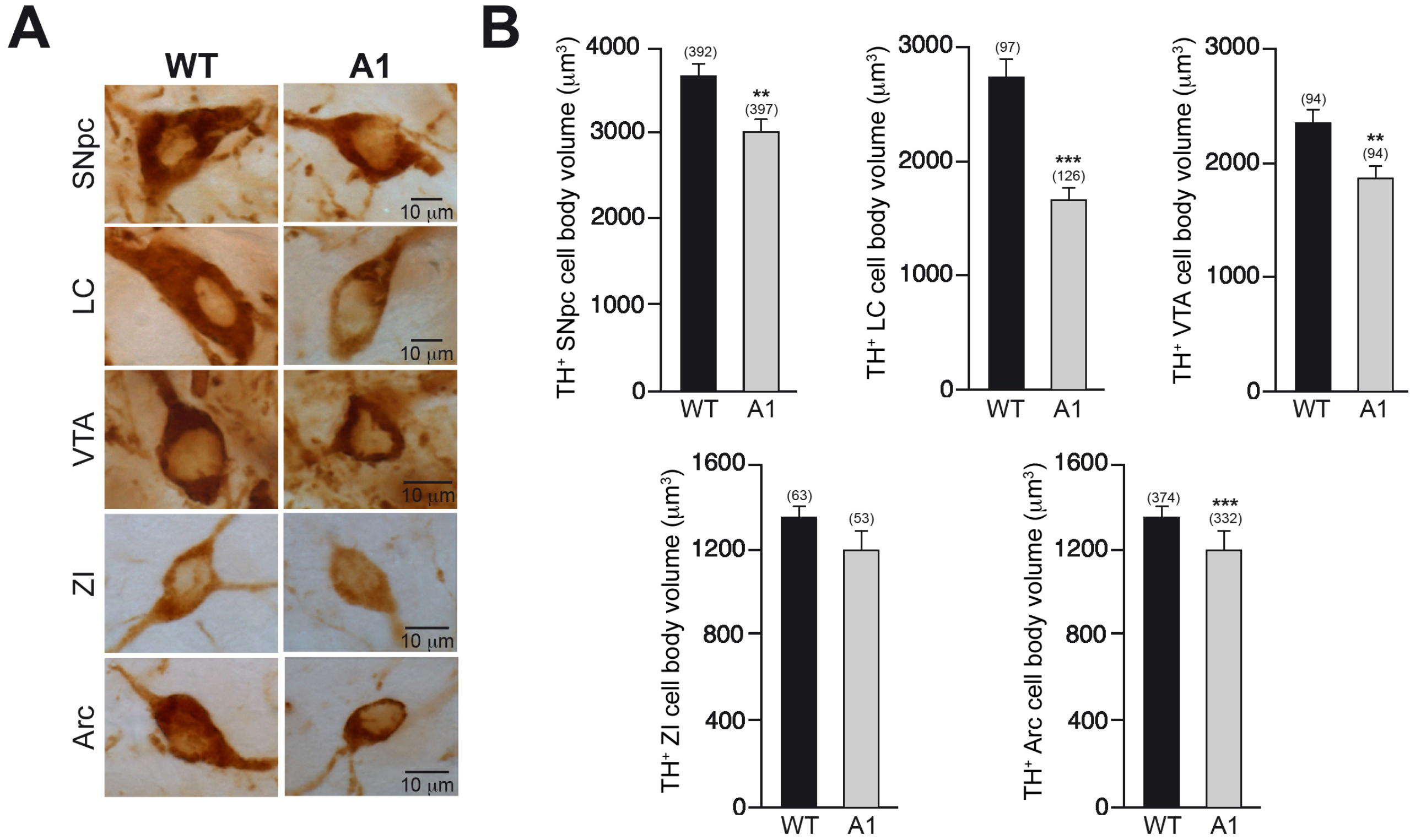
A1 overexpression decreases the cell body volume of CNS catecholaminergic neurons. A: High magnification images of representative TH^+^ neurons from different CNS catecholaminergic nuclei from wt and A1-mice. B: Stereological quantification of the neuronal cell body of catecholaminergic neurons from the nuclei and the experimental groups indicated. Data are presented as mean ± standard error of the mean (S.E.M.). The number of neurons analyzed are shown between brackets. Mann–Whitney U test (SNpc; LC; VTA and Arc); Unpaired t-test (ZI). **p<0.01; ***p<0.001.

### Structural alteration in the striatum of Anosmin-1 transgenic mice

The increased number of SNpc dopaminergic neurons found in the A1-mice prompted us to analyze their main target of projection, the striatum. The densitometric analysis of the TH^+^ striatal innervation showed only a minor increase in A1-mice in comparison with wt-controls (Fig. 3A,B). We analyzed in further detail the morphology of the striatum and observed that in A1-mice striosomes were significantly more numerous but smaller than in wt-controls (Fig. 3C-D). Moreover, when the area occupied by striosomes in respect to the striatal matrix was studied, the percentage of the striatal area occupied by striosomes was significantly larger in the A1-mice, with a subsequent decrease in the area occupied by the matrix (Fig. 3D).

**Figure 3.**
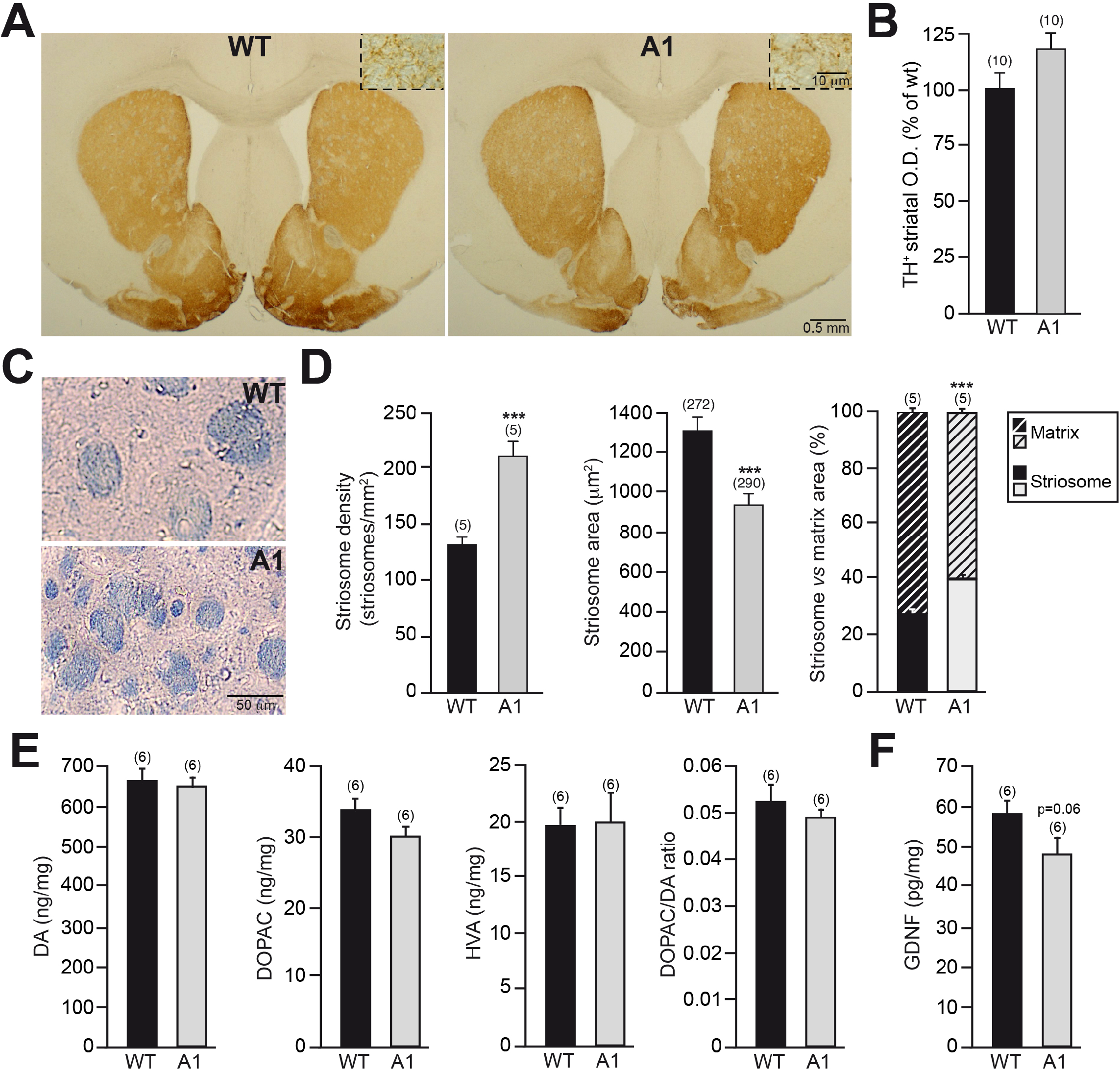
Altered striosome and matrix compartments in the striatum of A1-mice. A: Striatal coronal sections, after TH immunostaining, from wt and A1-mice. The insets depict, at high magnification, the dopaminergic striatal innervation. B: Analysis of striatal dopaminergic innervation by O.D. measurements. C: High magnification pictures, after eriochrome cyanine staining, showing the striosomes (highly myelinated) of wt and A1-mice. D: Stereological quantification of striosome density, area, and striosome *vs* matrix area. E: Striatal content of DA, DOPAC, HVA and DOPAC/DA ratio in wt and A1-mice. F: Striatal level of the dopaminotrophic factor GDNF in the wt and A1-mice. Data are presented as mean ± standard error of the mean (S.E.M.). The number of mice [B;D (striosome density and striosome *vs* matrix area); E;F] or striosomes [D, striosome area] analyzed are shown between brackets. Unpaired t-test [B;D (striosome density and striosome *vs* matrix area);E (DA, DOPAC and HVA content);F]; Mann–Whitney U test [D (striosome area); E, (DOPAC/DA ratio)]. ***p<0.001.

In addition to these striatal structural alterations, we investigated the neurochemical features of the striatum in A1-mice, without finding significant changes in the levels of DA and its metabolites DOPAC and HVA between A1-mice and wt-controls (Fig. 3E). Moreover, no differences were found when the ratio DOPAC/DA were examined between A1-mice and wt-controls, indicating a similar DA turnover between both experimental groups. Since GDNF has been shown to provide trophic support to the dopaminergic SNpc neurons and promote the arborization of TH^+^ terminals (Lin et al. 1993; Pascual et al. 2008; Enterría□Morales et al. 2020): A1-mice showed a trend of ∼20% reduction in the GDNF protein striatal levels respect to wt controls (Fig. 3F). Anosmin-1 overexpression has been previously reported to increase oligodendrogenesis and the number of mature oligodendrocytes in the corpus callosum (Murcia-Belmonte et al. 2016). Because A1-mice showed an important increase of the area covered by highly myelinated striosomes, we explored how this structural alteration could be related to an elevated oligodendrogenesis in the adult striatum. The stereological quantification of the density of mature striatal oligodendrocytes (CC1^+^ cells) in A1-mice and wt controls did not show any significant differences (Fig. 4A-B). Moreover, we found no significant difference in the density of striatal oligodendrocytes that were clearly located in the patch (striosome-CC1^+^ cells; Fig. 4C), while we detected a ∼10% reduction in the matrix compartment of A1 overexpressing mice (matrix-CC1^+^; cells Fig. 4D), in both cases we excluded those placed in the striosome/matrix interface. These results suggest that the augmented area covered by the patch compartment in A1-mice does not reflect higher oligodendrogenesis in the adult striatum.

**Figure 4.**
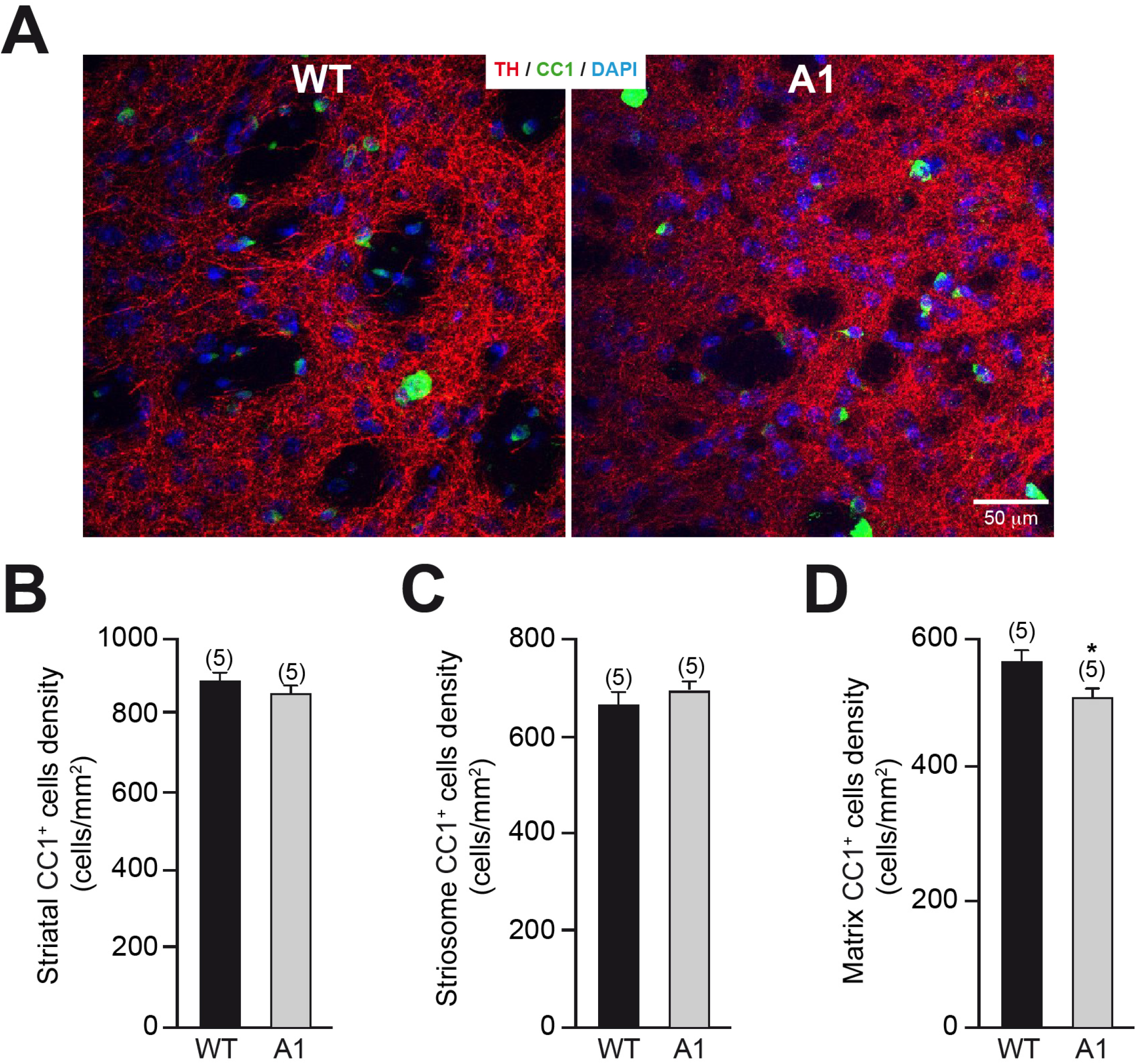
Analysis of the striatal oligodendrocytes in A1-mice. A: High resolution confocal images showing dopaminergic innervation (red) and CC1^+^ oligodendrocytes in the striatum from wt and A1-mice. B-D: Study of CC1^+^ oligodendrocyte density in the striatum (B) or located in the striosome (C) or matrix (D) compartment. In B-B, data are presented as mean ± standard error of the mean (S.E.M.). The number of mice analyzed are shown between brackets. Results of unpaired t-test are represented as: *p<0.05.

### Effects of Anosmin-1 overexpression in the nigrostriatal degeneration induced by chronic MPTP treatment

Considering the effects of A1 overexpression in the size and number of dopaminergic SNpc neurons and the structural organization of the striatum, we investigated how these morphological alterations of the nigrostriatal pathway could confer a differential susceptibility to chronic MPTP-induced parkinsonism. To do so, A1 and wt mice were treated for 3 months with MPTP, or vehicle solution, and the vulnerability of the nigrostriatal pathway was evaluated for each genotype. At the level of the striatum, the chronic MPTP treatment produced the degeneration of the dopaminergic terminals, measured as TH^+^ O.D., that was similar for both A1 and wt mice (Fig. 5A-B). The area of the striosome and matrix compartment was also analyzed in the striatum of the different experimental groups. As indicated in Figure 5C, wt mice treated with MPTP showed a slight increase in the striatal area occupied by the striosome compartment with respect to wt mice treated with saline. However, in the case of A1-mice, which present an important increase in the striatal area of the patch compartment, no further alterations were found associated to the chronic MPTP-treatment.

**Figure 5.**
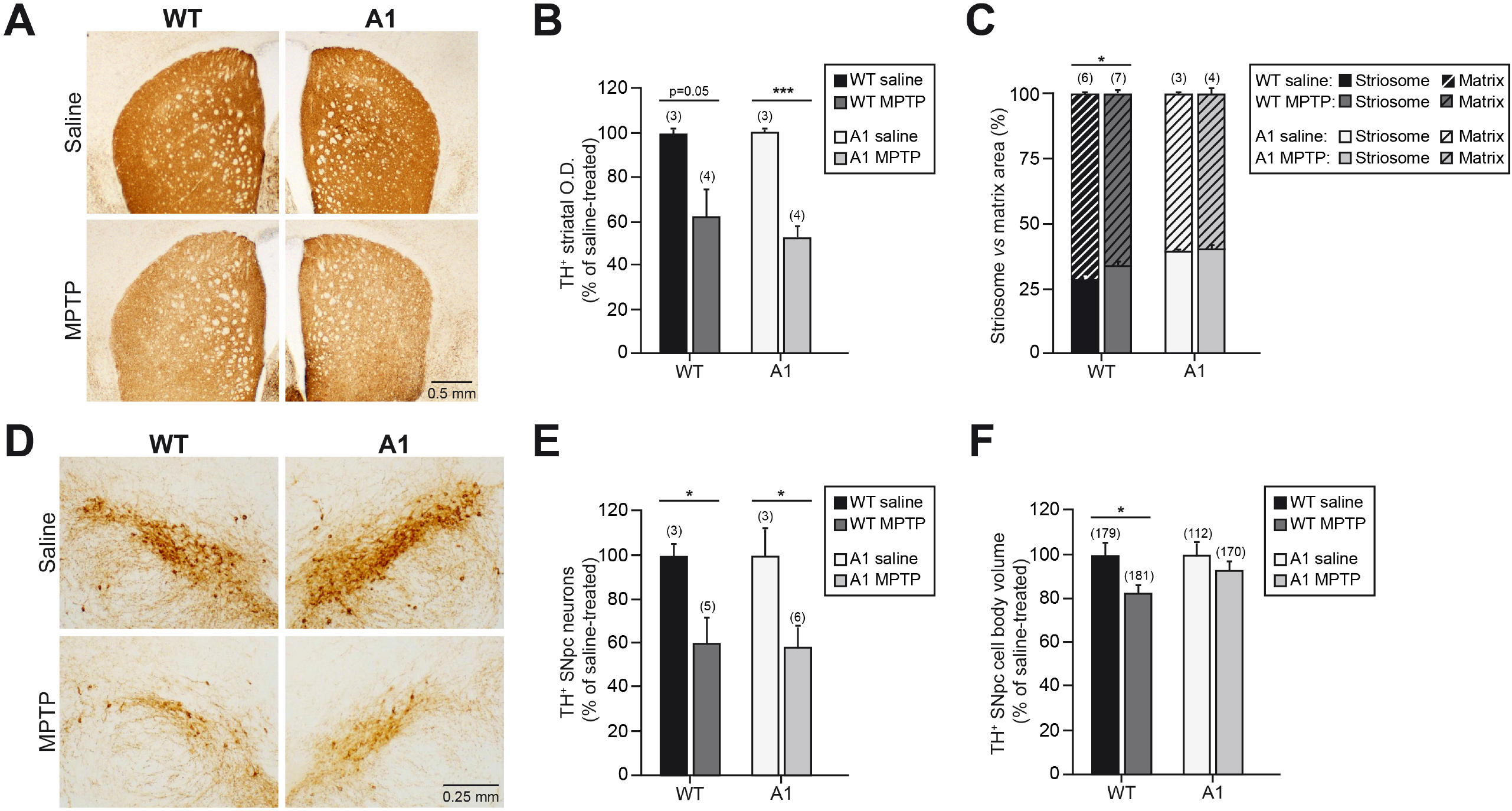
A1 overexpression does not alter the sensibility of the nigrostriatal pathway to chronic MPTP-induced neurodegeneration. A: Striatal coronal sections, after TH immunostaining, from wt and A1-mice chronically treated with vehicle (saline) or MPTP. B: Optical density (O.D.) analysis of the dopaminergic striatal innervation. C: Quantification of the striosome *vs* matrix area. D: Representative images of mesencephalic coronal sections showing the TH^+^ SNpc neurons from the aforementioned experimental groups. E-F: Analysis of TH^+^ SNpc neurons (E) and TH^+^ SNpc cell body volume. Data are presented as mean ± standard error of the mean (S.E.M.). The number of mice (B; C; E) or neurons (F) analyzed are shown between brackets. In B, E, and F data are presented as % of saline-treated mice of each genotype (wt or A1-mice) to reveal susceptibility to chronic MPTP-induced neurodegeneration. Results of unpaired *t*-test (B, C, E) and Mann–Whitney U test (F) are represented as: *p<0.05; ***p<0.001.

At the level of the SNpc, the TH^+^ cell death and the neuronal soma volume were measured as hallmarks of the neurodegenerative process induced by the experimental model of parkinsonism. As indicated in Figure 5D-E, the TH^+^ neuronal cell death induced by the chronic MPTP was similar for A1 and wt mice. The analysis of the TH^+^ neuronal soma volume in the remnant dopaminergic neurons showed a significant decrease (∼20%) in the neuronal soma volume in the wt mice subjected to chronic MPTP treatment. However, in the case of A1-mice, whose TH^+^ SNpc neurons are smaller than those from wt-mice as previously showed in figure 2, the MPTP treatment only produced a minor and not significant further reduction on the TH^+^ neuronal soma volume (Fig. 5F). Taken together, the comparative analyses performed in wt and A1-mice subjected to the chronic MPTP parkinsonian model indicate that despite the important structural alterations present in the nigrostriatal pathway of A1-mice, they exhibit a similar susceptibility to the MPTP-induced neurodegeneration than wt controls.

### Alterations in peripheral dopaminergic tissues induced by Anosmin-1 overexpression

To further investigate the effects of A1 overexpression in the PNS, we analyzed the morphological features of different dopaminergic tissues like the superior cervical ganglia (SCG), the carotid bodies (CB) and the adrenal medulla (AM). The histological analysis of the sympathetic SCG did not show any difference in the anatomical location, the general organ appearance or volume when organs obtained from A1-mice were compared to those extracted from wt controls (Fig. 6A-B). However, despite the localization of the adjacent CB being also unaffected, the morphological analysis of the CBs from A1-mice revealed a significant reduction in the organ volume when compared to wt CBs (Fig 6C-D). Interestingly, that decrease in the CB volume of A1-mice corresponds with a diminished number of the neuron-like TH^+^ CB glomus cells (Fig. 6D). Apart from TH^+^ glomus cells, the CB is also composed of glia-like GFAP^+^ sustentacular cells (López-Barneo et al. 2009), which have been demonstrated to act as neural stem cells promoting the CB growth in hypoxia condition and participating in the organ postnatal maturation (Pardal et al. 2007; Díaz□Castro et al. 2015). Although the number of GFAP^+^ CB cells was only slightly reduced in A1-mice respect to wt controls, the GFAP^+^ O.D. analysis revealed a decreased GFAP expression in sustentacular cells from A1-mice (Fig. 6E,F), suggesting that A1 overexpression could alter the functionality of these glial cells.

**Figure 6.**
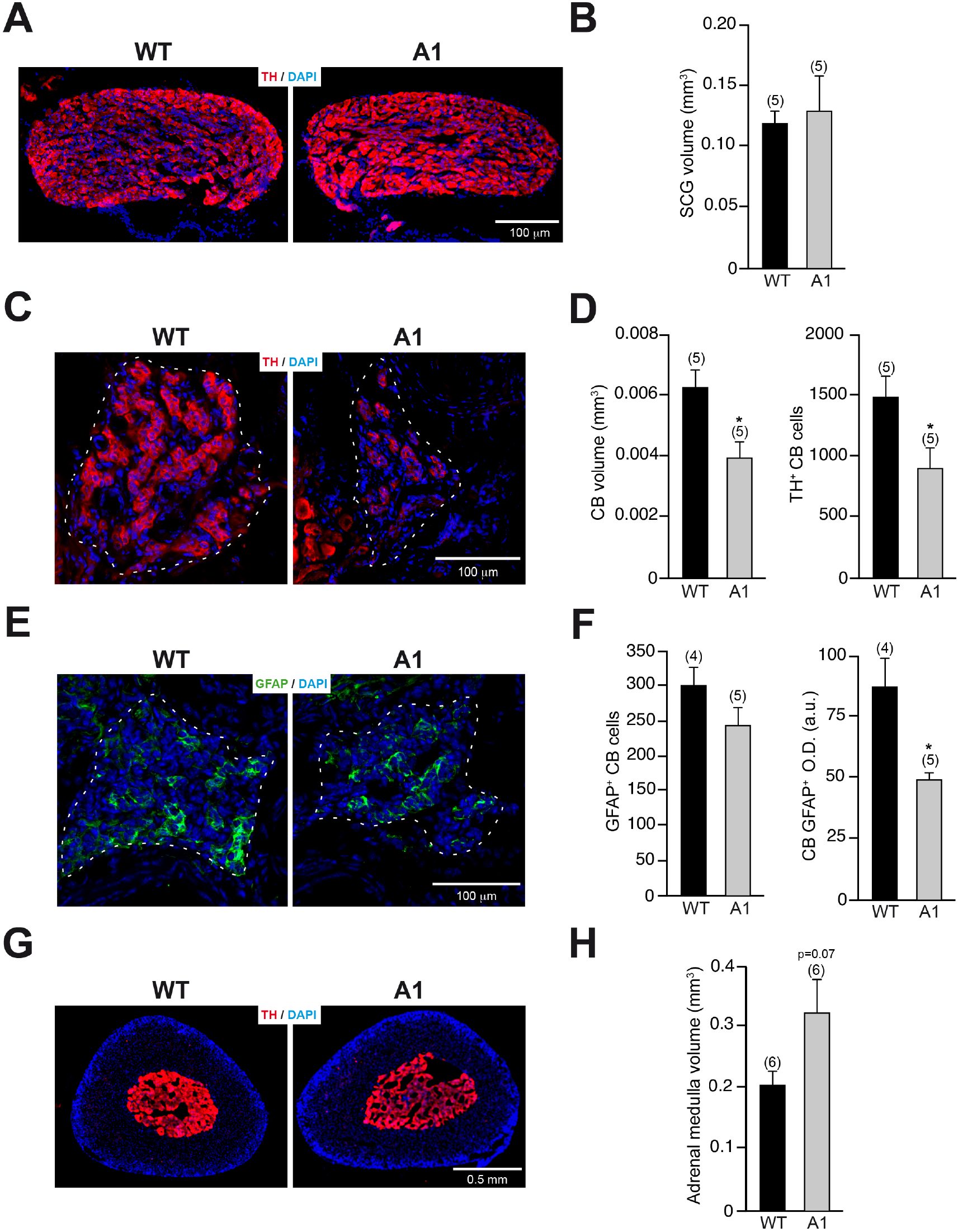
Morphological analysis of dopaminergic tissues associated with the PNS in A1-mice. A: Representative images, after TH immunofluorescence, of SCG from wt and A1-mice. B: Quantification of the SCG volume. C: Histological sections of CBs from wt and A1-mice, after TH immunofluorescence, showing chemosensitive TH^+^ CB glomus cells. D: Quantification of the CB volume and number of TH^+^ glomus cells. E: Histological sections of CBs from the previously described groups, after GFAP immunofluorescence, revealing GFAP^+^ CB sustentacular cells. F: Quantification of the number of GFAP^+^ CB sustentacular cells and the level of GFAP expression, by GFAP^+^ O.D. G. Representative images, after TH immunostaining, of suprarenal gland of wt and A1-mice. H: Quantification of the adrenal medulla volume. Data are presented as mean ± standard error of the mean (S.E.M.). The number of mice analyzed are shown between brackets. Results of unpaired t-test are represented as: *p<0.05.

Finally, the effect of A1 overexpression in the catecholaminergic AM was also studied. Suprarenal glands from A1-mice showed similar weight (3.97±0.31 mg for wt mice *vs*. 3.41±0.21 mg for A1-mice). However, the quantification of the TH^+^ AM volume between A1 and wt-mice showed a trend to increase the volume occupied by the catecholaminergic AM in A1-overexpressing mice (Fig. 6G-H). Taken together, the analysis of the effects produced by A1 overexpression in the three different catecholaminergic PNS tissues analyzed (SCG, CB and AM) revealed distinctive outcomes on the different tissues, varying from the lack of alterations in SCG to the decrease or increase in the TH^+^ volume of CB or AM respectively. These differential effects of A1 overexpression could suggest specifics niche-dependent actions of A1 in the genesis and/or maturation of peripheral nervous system dopaminergic cells.

## DISCUSSION

Most of the biological effects of A1 have been investigated during neural development, but there are very few reports on its action on the adult mammalian nervous system (García-González et al. 2016; Murcia-Belmonte et al. 2016). Our present work extends the study of the effects of A1 to other known populations of catecholaminergic neurons in the adult CNS and PNS of the mouse. One of the novelties of the present report is that A1 overexpression produces larger populations of TH^+^ neurons (∼30-50% neurons more than in wt-control mice) in two main catecholaminergic structures of the CNS, as SNpc and LC. Interestingly, this increased number of TH^+^ neurons showed significantly smaller somas (∼10-40% smaller, depending on the nucleus studied). In closer relationship with our data, it has been reported that the absence of FGF-2 results in a larger number of dopaminergic neurons in the SNpc and the OB (Ratzka et al. 2012; Vergaño-Vera et al. 2015). Among other components of the extracellular matrix, such as heparan-sulfates, perlecan or laminin, A1 has been shown to regulate FGF availability and biological activity through different ways, including FGFR1 interaction and shuttling (Bribián et al. 2006, 2020; Clemente et al. 2011; Endo et al. 2012; de Castro et al. 2014, 2017; Díaz-Balzac et al. 2015; Wang et al. 2018). Our present study would reflect more evidence in this sense, confirming that FGFR1 signaling may be important for neurogenesis and/or neuronal differentiation toward dopaminergic phenotypes, in agreement with the previously reported depletion of dopaminergic neurons when a dominant negative FGFR1 is expressed under the control of the TH promoter (Klejbor et al. 2006).

The examination of the dopaminergic striatal innervation revealed only a non-statistically significant slight increase in A1-mice. However, the joint analysis of the number of dopaminergic SNpc neurons (∼30% of increase), the dopaminergic striatal innervation (∼15% of increase) and the striosome/matrix area observed in A1-mice suggest that A1 overexpression induces a defect on the striatal arborization of SNpc dopaminergic neurons. This could be due to more numerous but smaller TH^+^ SNpc neurons giving rise to a relatively less ramified striatal fibers, which also occupies a smaller striatal area. This diminished striatal fiber sprouting of nigrostriatal SNpc neurons could be related to the trend of ∼20% reduction in the striatal GDNF content observed in A1-mice, since it is well known that this neurotrophic factor promotes the fiber sprouting of dopaminergic fibers in the striatum (Lin et al. 1993; Airaksinen and Saarma 2002).

Another relevant finding of our study is the structural alterations observed in the striatum of A1-mice, which present more numerous yet smaller striosomes than in wt-controls. Our data indicate that this alteration does not reflect significant changes in oligodendrogenesis in the adult striatum, which contrasts with the effects observed in the main brain commissure (Murcia-Belmonte et al. 2016), and confirms the great degree of heterogeneity displayed by oligodendroglia, in general, and oligodendrocyte precursors, in particular (Jäkel et al. 2019; Bribián et al. 2020; Chamling et al. 2021; Yaqubi et al. 2022). During embryonic development striosomes are established and differentiated in the “dopamine islands” constituted by the incoming dopaminergic nigrostriatal fibers (Crittenden and Graybiel 2011). The fact that A1-mice present significantly higher number of dopaminergic SNpc neurons projecting to the striatum supports the hypothesis that the striosome/matrix imbalance observed in these mice could be related to the altered nigrostriatal dopaminergic input produced by A1 overexpression.

A recent study has revealed an interesting relationship between the nanoscale organization and diffusion of extracellular matrix components and neuronal cell death in a model of α-synuclein induced neurodegeneration (Soria et al. 2020). In addition, an increased number of dopaminergic neurons in the olfactory bulb of parkinsonian patients has been reported (Mundiñano et al. 2011), suggesting the possibility that increased dopaminergic tone of certain nuclei could alter the susceptibility to neurodegeneration. In this work, we studied how the morphological changes observed in the nigrostriatal pathway of A1-mice (with more and smaller dopaminergic SNpc neurons and altered patch/matrix ratio in the striatum) could confer a differential susceptibility to experimental parkinsonism, but our experiments revealed that A1-mice show similar susceptibility to the MPTP-induced neurodegeneration than wt controls. Our present results have relevant pathogenic implications, because after them we would discard, at least in our chronic MPTP model, the relevance of the initial number of dopaminergic neurons in SNpc and/or LC and the striatal patch/matrix ratio in the pathogenesis of PD.

Regarding the effect of A1 overexpression in the PNS, of the three catecholaminergic tissues analyzed (SCG, CB and AM), only the CB shows evident morphological changes, with a clear reduction in the organ volume and the number of the chemosensitive TH^+^ glomus cells. Interestingly, CBs from A1-mice also show histological alterations in GFAP^+^ sustentacular cells that have been identified as neural stem cells in the adult organ, promoting the postnatal growth and maturation (Pardal et al. 2007; López-Barneo et al. 2009; Díaz□Castro et al. 2015). In relation to the role of A1 as a main regulator of FGF availability, it has been previously demonstrated that FGF2 promotes the proliferation and survival of CB TH^+^ glomus cells (Nurse and Vollmer 1997), and the absence of docking protein FRS2α, an important mediator of the FGF signaling pathway, induces the agenesia of this organ (Kameda et al. 2008). Therefore, our present data suggest that FGF signaling pathway may have a relevant role in the CB neurogenesis and A1 modulates this, as shown in neural crest early development (Endo et al. 2012).

Our work confirms previously reported data (García-González et al. 2016) and indicates that A1 plays a relevant role in the specification and survival of catecholaminergic neurons in different neuronal nuclei of the mammalian nervous system. The different effects of A1 overexpression found in the catecholaminergic nuclei analyzed, strongly suggest specific niche-dependent actions of A1 in the development of these neurons. Strikingly, A1 overexpression induces, in terms of TH^+^ neuronal number, opposite effects between the CNS (increasing the number of TH^+^ neurons in OB, SNpc and LC) and the PNS (reducing the number of TH^+^ neurons in the CB). However, this opposed effects on the genesis of dopaminergic neurons between the CNS vs PNS seems to be mediated, in both cases, through an abortive interaction with the FGF signaling pathway, since the *in vivo* knock down of main components of this pathway produces a phenotype similar to A1 overexpression both in SNpc (Ratzka et al. 2012; Hövel et al. 2019) or in CB (Kameda et al. 2008). In conclusion, A1 appears as a principal regulator of the FGF signaling pathway in relation to the formation, differentiation, and survival of catecholaminergic neurons of different nuclei of the mammalian nervous system.

## Declarations

### Ethics approval and consent to participate

Animal procedures were approved by the Animal Review Board at the Hospital Nacional de Parapléjicos (registered agreement number SAPA001), Instituto Cajal-CSIC (CEEA-IC) and the Animal Research Committee of the University Hospital Virgen del Rocío (University of Seville; registered agreement number 11-07-14-112).

### Availability of data and material

All relevant data are included in the paper. This study did not generate data sets deposited in external repositories. Information/data required will be available by the corresponding authors upon request.

### Competing interests

The authors have no relevant financial or non-financial interests to disclose.

### Funding

This research was supported by MCIN/Spanish Research Agency (AEI)/ 10.13039/501100011033 grants PID2019-105995RB-I00 (J.J.T.-A. and J.V.) and PID2019-109858RB-I00 (to F.d.C.); ISCIII grants Red TerCel, RD16/0011/0025 (J.J.T.-A.), PI12/02574 (J.J.T.-A.), CP21/00106 and PI22/00156 (DG-G); Consejería de Innovación, Ciencia y Empresa grant CTS2739 (J.J.T.-A.), Consejería de Economía, Conocimiento, Empresas y Universidad grant US-1380891 (J.J.T.-A. and J.V.), Junta de Andalucia; grant 2019AEP033 from Consejo Superior de investigaciones Científicas-CSIC (to F.d.C.); Proyectos de Investigación Relacionados con las Enfermedades Raras Infantiles (to F.d.C.), Fundación Inocente-Inocente (Spain). C.N. is a Tomás y Valiente fellow (MIAS-UAM).

### Authors’ contributions

Conceptualization: J.V.; F.dC. and J.J.T.-A. Funding acquisition: J.V.; F.dC. and J.J.T.-A. Methodology and investigation: J.V.; R.G.-S.; F. dC.; D.G.-G.; R.L.-G.; E.G.-R.; N.S.-L.; C.N.; M.M. Data analysis: J.V.; R.G.-S.; E.G.-R. Project supervision: J.V.; F.dC. and J.J.T.-A. Visualization: J.V.; R.G.-S.; E.G.-R. Writing—original draft: J.V.; F.dC. and J.J.T.-A; Writing—review and editing: J.V.; F.dC. and J.J.T.-A. All authors have read and agreed to the published version of the manuscript.

## Acknowledgements

The authors thank to A. Bermejo-Navas for excellent technical help.

## Notes

### Competing Interest Statement

The authors have declared no competing interest.

